# Dynamic Control of Ca^2+^ Binding in the C2 Domains of Synaptotagmin 1

**DOI:** 10.1101/412031

**Authors:** Patrick J. Rock, Austin G. Meyer, Chantell S. Evans, Edwin R. Chapman, R. Bryan Sutton

**Affiliations:** Texas Tech University Health Sciences Center, School of Medicine, 3601 4th Street, 79430 Lubbock, TX, USA; Ohio State University Medical Center, Departments of Internal Medicine and Pediatrics, 410 West 10th Avenue, 43210 Columbus, OH, USA; Texas Tech University Health Sciences Center, Department of Cell Physiology and Molecular Biophysics, Center for Membrane Protein Research, 3601 4th Street, 79430, Lubbock, TX, USA; University of Wisconsin, Howard Hughes Medical Institute and Department of Neuroscience, 1111 Highland Avenue, 53705, Madison, WI, USA

**Keywords:** synaptotagmin, C2 domain, Ca^2+^ binding protein, AD3 mutation, exocytosis

## Abstract

Synaptotagmin senses fluctuations in the Ca^2+^ environment of neurons near active zones and transduces a signal to the SNARE complex to initiate exocytosis at the presynaptic terminus. The 3D structures of the two tandem C2 domains of synaptotagmin have been determined to high resolution; however, it is currently unclear how each domain dynamically interacts with Ca^2+^ at the atomic level. To study the mechanistic consequences of the lethal mutations at the AD3 locus, we introduced tyrosine to asparagine point mutations in both the C2A and C2B domains of synaptotagmin 1, and we have constructed a model that describes the relationship between Ca^2+^ -binding and the structural changes within each C2 domain. We show that the mobility of loop 3 in the Ca^2+^ binding pocket increases markedly in C2A, while the mobility of loop 1 changes in C2B with the AD3 mutation. This increase in loop mobility results in an increase in the average volume and variance of the Ca^2+^ -binding pockets of C2A and C2B. The volume of the unbound Ca^2+^ -binding pocket in C2A is usually restrained by intra-domain interactions between the tyrosine residue at the AD3 locus and residues on loop 3; however, the AD3 mutation decouples the restraint and results in a larger, more variable Ca^2+^ -binding pocket in C2A. C2B maintains a more compact Ca^2+^ -binding pocket; however, its volume also fluctuates significantly with the AD3 mutation. Changes in binding pocket volume that involve more variable Ca^2+^ binding loops would likely affect Ca^2+^ affinity in the neurons of the affected organism. Using molecular-dynamics simulations, we show that mutations at the AD3 locus alter the mobility of the Ca^2+^ -binding loops by removing a key stabilization mechanism that is normally present in C2 domains. The lack of loop stabilization results in a net increase in the volume of the Ca^2+^ -binding pocket and provides an explanation for the observed lethal phenotype.

## Background

Exocytosis in neurons occurs with high temporal and spatial fidelity. The fusion of neurotransmitter-containing vesicles with the presynaptic terminus occurs only at active zones on the nerve terminal under the influence of a tightly controlled Ca^2+^ gradient. Synaptotagmin 1 (Syt1) is now accepted as the Ca^2+^ sensor for synchronous neuronal exocytosis [1,2], whereas the formation of the SNARE complex provides the free energy to fuse the phospholipid bilayers of the vesicle and the presynaptic terminus [3]. Ca^2+^, synaptotagmin, and the SNARE complex, therefore, comprise the minimum components necessary to fuse membranes in a Ca^2+^ - dependent manner [4,5,6]. There are at least 16 paralogs of synaptotagmin that have been annotated in the human genome [7]. Typically full-length synaptotagmin proteins begin with a short intravesicular peptide at the amino-terminus, followed by a single *α*-helical transmembrane span, a variable length tether, and two homologous, tandem C2 domains designated as C2A and C2B [8,9]. The two C2 domains are joined by a short linker that varies in size depending on the isoform [10,11,7]. Each C2 domain is an independently folded Ca^2+^ -dependent phospholipid-binding domain composed of two *β*-sheets, each made up of four *β*-strands [12,13]. The *β*-sheets are curved under the influence of multiple *β*-bulges, some of which are unique to C2 domains [14]. In both C2 domains of Syt1, calcium ion is coordinated by five conserved Asp residues located on loops 1 and loop 3 at the apex of the fold. Loop 2 delineates one side of the Ca^2+^ binding pocket, but does not contribute coordinating residues. In addition loops 1 and 3 directly participate in the lipid anchoring activity of the domain through two hydrophobic residues at the tips of the loops 1 and 3 [15,16,17]. A considerable number of X-ray and NMR structures of isolated C2 domains of synaptotagmin are now available; however, far fewer tandem C2 domain structures have been determined [10,18,19,11].

Early in the scientific debate to clarify the role of synaptotagmin in exocytosis, a point mutation (AD3) within the *Drosophila* synaptotagmin gene was generated that resulted in a lethal phenotype [20]. The mutation was characterized by sluggish crawling and defective feeding behavior in early instars. Further, the larvae did not grow or molt and died after about one week [20]. In the neurons of these animals, the overall amounts of the protein that localized to nerve terminals were like that of wild-type synaptotagmin, so protein misfolding or aberrant protein localization were excluded as the explanation of this phenotype [21,22]. At low Ca^2+^ concentrations, evoked release was essentially abrogated in AD3 mutants [23]. Synaptotagmin carrying the AD3 mutation was found to bind SNAREs *in vivo*; therefore, despite its lethal phenotype, the stability of the mutated domain was likely similar to wild-type [22,23]. Further study of this mutation showed that it was possible to rescue the lethal phenotype by exposing the fly larvae to higher concentrations of extracellular Ca^2+^ [23]. Sequencing of the mutated gene revealed that a tyrosine at position 364 was changed to an asparagine in the C2B domain of the *Drosophila* synaptotagmin gene [22]. From a physiological perspective, the AD3 mutation still functioned, albeit with a severe deficit. Interestingly, the affected tyrosine residue is part of a conserved sequence motif present in most C2 domains, not only in synaptotagmin (Fig 1) [24]. Given the location of this mutation on *β*- strand 3 in C2B, it is unclear how it could affect Ca^2+^ sensitivity; yet, it is clear that there is a Ca^2+^ binding defect in the *Drosophila* Syt1 C2B domain [25].

**Figure 1.**
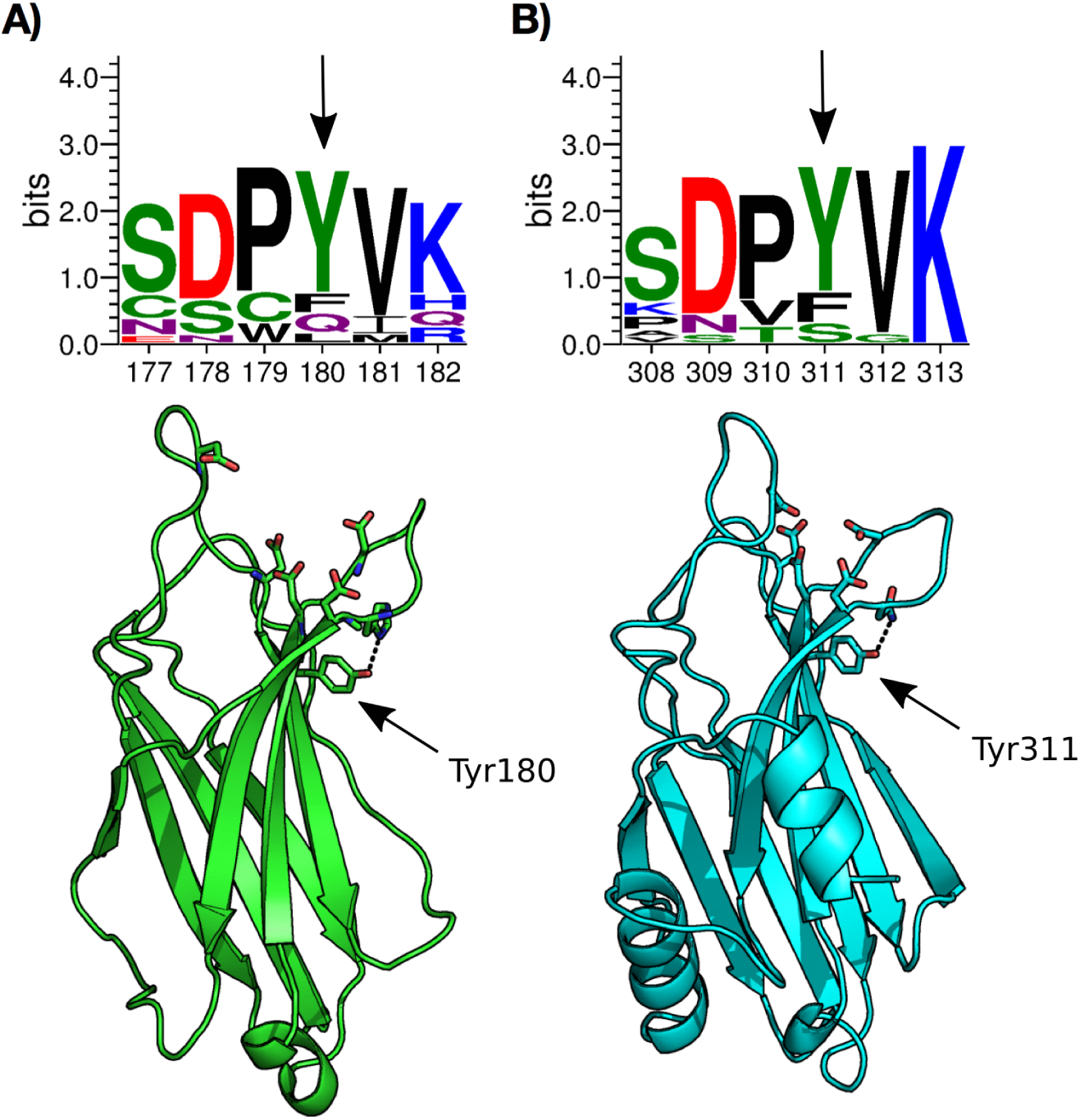
Conservation and location of the AD3 mutation in C2A and C2B. Syt C2A and C2B sequence logos extracted from an alignment of 16 human synaptotagmin isoforms. In A, C2A is displayed. The sequence is truncated to include only the C2A residues 176-182 (human Syt1 numbering) that juxtapose the AD3 locus. In B, C2B is displayed. The sequence is truncated to include only C2B residues 308-313 that juxtapose the AD3 locus. The mutation site is highlighted on the lower half of each sub-figure. The relative font size of each residue indicates the overall sequence conservation of that amino acid at that position in the entire alignment.

The available genetic and physiological data suggest that mutations at AD3 disrupt a link between the C2 domain structure and the domain’s ability to respond to the dynamic Ca^2+^ environment that exists at the active zone. To understand this link in greater detail, we studied the impact of the AD3 mutation in the isolated C2A and C2B domains of Syt1 using molecular dynamics simulations. We show that AD3 mutations in both C2 domains result in markedly increased mobility of the Ca^2+^ -binding loops and that this is most likely due to a loss of intra-domain contacts. In our simulations, the contacts appear to be mediated by the conserved tyrosine residue at the AD3 locus in each domain. Interestingly, the Ca^2+^ -binding loops of C2A and C2B are affected differently by AD3 mutations. C2A loop 3 is made more flexible after the AD3 mutation. By contrast loop 1 is made more flexible by the AD3 mutation in C2B. The AD3 locus, therefore, is a critical feature of the Ca^2+^ -binding machinery that restricts loop mobility and stabilizes the Ca^2+^ - binding architecture within C2 domains.

## Results

### Minimal SNARE-based Fusion Assay

To test the Ca^2+^ -dependence of Syt1 in the context of the original Syt1 AD3 mutation (Y311N), we used two variants of an *in vitro* SNARE-based fusion assay to compare the ability of the cytoplasmic domain of wild-type (C2AB) and the AD3 mutant (C2AB AD3) to mediate Ca^2+^ -dependent vesicle fusion (Fig 2). In the standard fusion assay, we measured the percentage of *in vitro* vesicle fusion as a function of time (Fig 2A and 2C). C2AB failed to facilitate vesicle fusion in the absence of Ca^2+^ (EGTA curve) and achieved ~50% maximal fusion in the presence of 1 mM Ca^2+^. However, C2AB AD3 triggered less Ca^2+^ -dependent fusion (*∼*30% maximum fusion) under similar experimental conditions. In the split t-SNARE variation of the fusion assay, the ability of Syt1 to fold soluble SNAP-25 onto syntaxin was also disrupted by the mutation in a Ca^2+^ -dependent manner (Fig 2B and 2C) [26]. Compared to the standard fusion assay where the SNARE complex was preformed prior to synaptotagmin engagement, there was a marked decrease in fusion using the C2AB AD3 (Fig 2B and 2C). Therefore, the AD3 mutation in C2B, which was not directly associated with the Ca^2+^ -binding pocket, showed a defect in the Ca^2+^ -dependent vesicle fusion in this minimal component assay. To understand how the AD3 locus interacts with the calcium ion binding residues of the C2 domains of Syt1, we performed a series of molecular-dynamics simulations on wild-type and mutant Syt1 C2 domains, with and without Ca^2+^.

**Figure 2.**
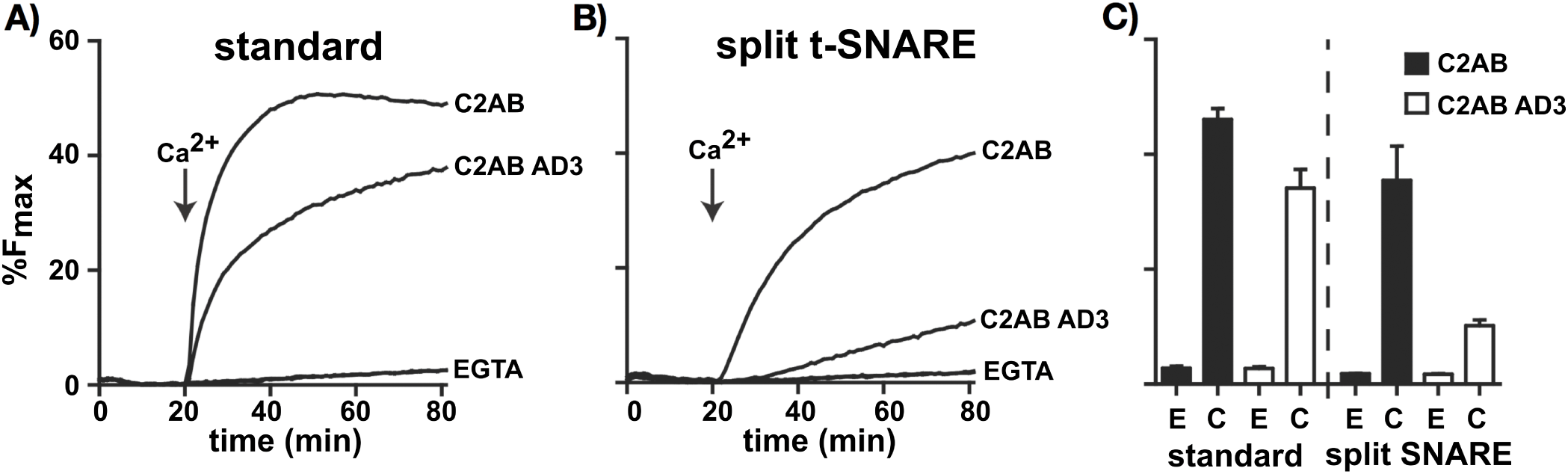
The AD3 mutant does not stimulate fusion as efficiently as Syt1. The cytoplasmic domains (C2AB) of Syt1 and the AD3 mutant were examined using two forms of the *in vitro* membrane fusion assays. In A, proteins were initially screened in the standard *in vitro* fusion assay; both proteins stimulated fusion with the AD3 mutation functioning at a slight deficit. In B, these proteins were examined in a more stringent form of the traditional standard assay. In the split t-SNARE assay, Syt1 must fold SNAP25B onto Syx1A to stimulate fusion. The AD3 mutant failed to efficiently stimulate fusion compared to Syt1. For comparison, representative traces of Syt1 and the AD3 mutant for both fusion assays are provided. (C) Bar graph summarizing the extent of fusion (t = 80 min). ‘E’ represents the assay with EGTA, and ‘C’ represents the assay with Ca^2+^. Data presented as mean *±* SEM; n = 3.

### Dynamic comparison of wild-type and mutant Syt1 C2 domains

To study the dynamic changes that occurred between wild-type and mutant C2 domains of Syt1, we compared the RMSF (root-mean-squared-fluctuations) of each C*α* without Ca^2+^ (Fig 3A and 3B). While there was little overall difference noted between the various simulations of the wild-type and mutant C2 domain, the loops of the binding pocket showed considerable variance. The AD3 mutation yielded the greatest difference at loops 1, 2, and 3 in the mutant forms of both the C2A and C2B domains, Y180N (C2A^AD3^) and Y311N (C2B^AD3^), respectively (Fig 3A and 3B). The most significant difference as a result of the mutation was the doubling of the RMSF in loop 1 of C2B. We also measured a smaller increase in RMSF in loop 2 of C2B (Fig 3B). C2A showed less pronounced RMSF differences between C2A^AD3^ and C2A^WT^, but there was a marked increase of the RMSF in loop 3 (Fig 3A). Repeating the analyses with two calcium ions occupying the binding sites of each C2 domain showed similar RMSF differences between mutant and wild-type across the entire protein domain. The reduction of the amplitude of the RMSF values of the loop residues in the Ca^2+^ -bound state was likely a function of the stabilization of the domain imparted by the interactions between calcium ion and acidic residues on each loop. The net result was that calcium ions bridge loops 1 and 3, thus reducing the overall flexibility of the loops. In a homologous system, Ca^2+^ -dependent restriction of loop mobility was also reported for the single C2 domain of protein kinase C using molecular-dynamics simulations [27] and NMR techniques [28].

**Figure 3.**
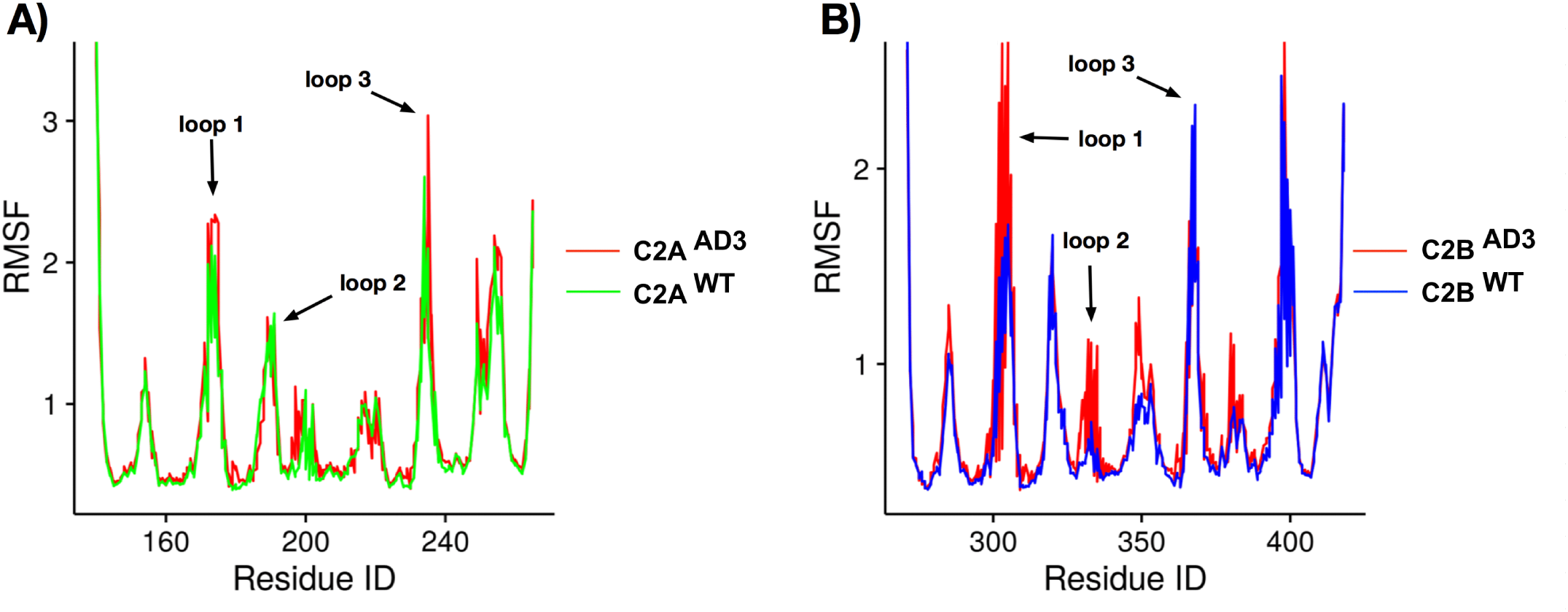
RMSF comparison by residue of C2A and C2B with and without the AD3 mutation. In A, we show RMSF of C2A domain C*α* in Ca^2+^ unbound state. The green line shows the wild-type and the red line shows the AD3 mutant. In B, we show RMSF of C2B domain C*α* in Ca^2+^ unbound state. The blue line shows the wild-type. The red line shows the AD3 mutant. In both subfigures, the loops that form the bounds of the Ca^2+^-binding pocket are highlighted.

### Iso-Dynamic Comparison of Loop Flexibility

To assess the relative flexibility of each of the residues in the mutated C2 domains, we calculated the RMSF of each of the C*α*s after concatenation and alignment of the molecular dynamics trajectories. We then computed the difference in RMSF between the wild-type and AD3 mutant by correlating the RMSF values from each set of trajectories against one another. We plotted wild-type RMSF on the x-axis and AD3 mutant RMSF on the y-axis for each of the residues in the loops. The resulting coordinates represented the relative change in loop residue flexibility upon mutation (Fig 4A and 4B left). In this representation, if the correlation between wild-type and AD3 mutant falls on the iso-dynamic line (i.e. the identity or *x* = *y* line), there is no significant difference in mobility of that particular residue. Points above the iso-dynamic line denote higher mobility in the mutant; whereas, points below the diagonal line denote lower mobility in the mutant. To visualize these differences on the protein structure, we measured the residuals by calculating the vertical distance from each point to the iso-dynamic diagonal line. Positive residuals indicate greater mobility in the AD3 mutant for that particular residue, whereas negative residuals indicate lower mobility for the mutant. We then mapped the residuals onto a ‘putty’ representation of the C2 domains using Pymol ([29] (Fig 4A and 4B right). We found little difference in the overall dynamics of the protein within the body of the domain as illustrated by the width of the tubes in the cartoon representation; however, we found clear differences in the dynamic properties of loops 1, 2 and 3 in the unbound state (Fig 4A and 4B). Without Ca^2+^ to bridge loops 1 and 3 in C2A^WT^ and C2A^AD3^, the most significantly correlated points occurred above the diagonal in loop 3, indicating a greater mobility of that loop (Fig 4A). Interestingly, loop 3 in both C2B^WT^ and C2B^AD3^ was relatively rigid throughout the trajectory (Fig 4B), and loop 1 was more mobile.

**Figure 4.**
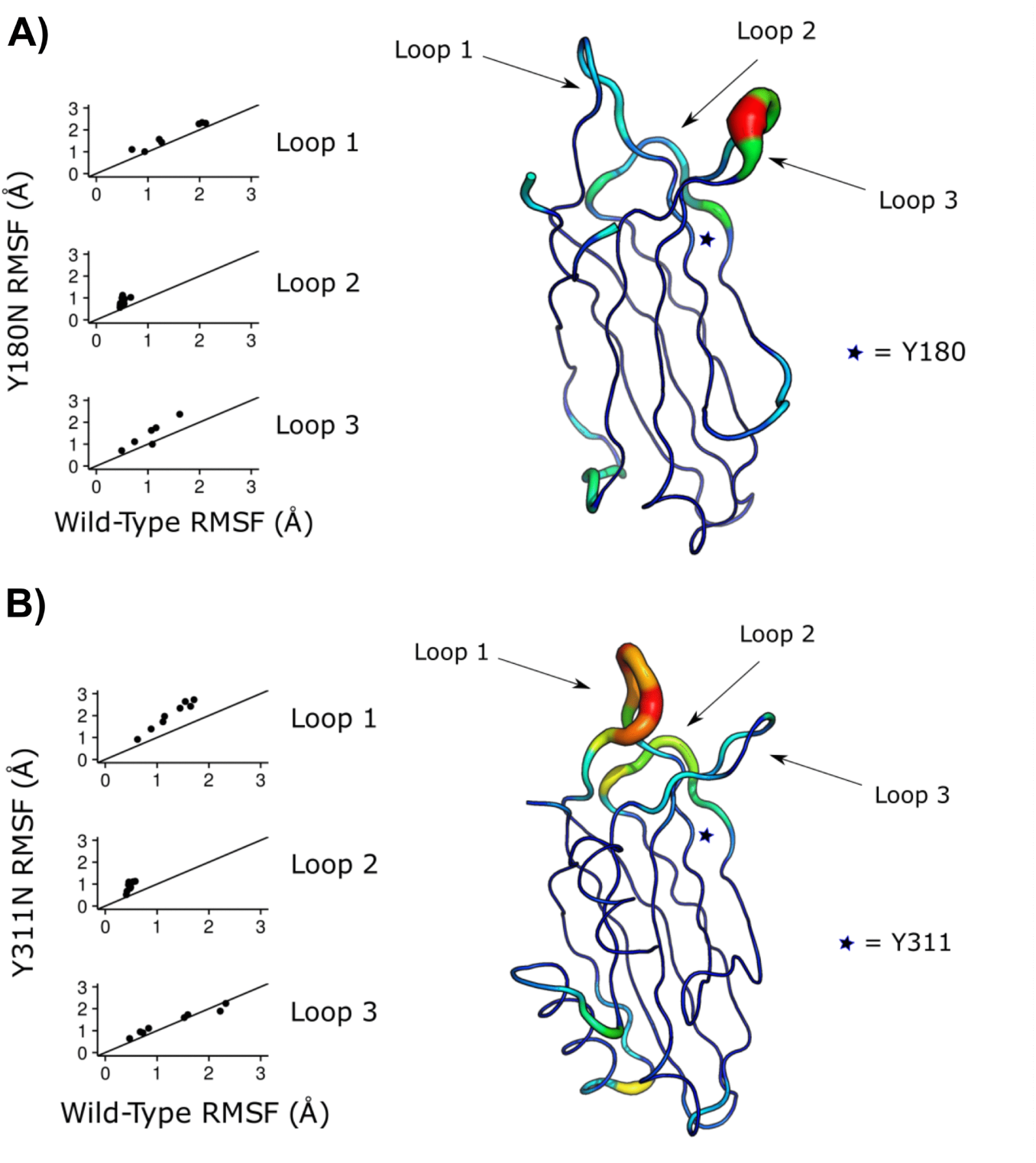
Comparative Iso-Dynamic Analysis of C2A and C2B. Dynamics analysis of C2A and C2B upon mutation at AD3. Each point is a particular residue in the loop indicated. Points above or below the line show changes with calcium bound while points on the line show no change. In A, on the left we show the iso-dynamic comparison of Ca^2+^ binding loop residues in C2A without Ca^2+^. On the right, we show the differential motion analysis plotted onto the structure. Sites that are thicker and more red show a greater difference between mutant and wild-type. The star represents the site of the AD3 mutation. In B, we show the iso-dynamic comparison of Ca^2+^ binding loop residues in C2B without Ca^2+^. The star represents the site of the AD3 locus. Mutation at the AD3 locus in C2A causes significantly increased mobility in loop 3. Mutation at the AD3 locus in C2B causes significantly increased mobility in loop 1 and to a lesser extent loop 2.

### Changes in the Ca^2+^ binding pocket

In many C2 domains, calcium ions are coordinated by oxygen atoms from a combination of four or five conserved acidic residues and backbone carbonyl oxygens. Five oxygen atoms from within each C2 domain of Syt1 coordinate Ca^2+^ with hexadentate geometry within a cup-shaped binding pocket. The sixth solvent-exposed axial coordinate is thought to be contributed by a phospholipid headgroup, thereby providing a link between the C2 domain protein component, Ca^2+^ occupancy, and phospholipid [24,30]. Since AD3 alters the mobility of the primary loops that comprise the divalent-cation binding pocket (Fig 3), we measured the variability of the binding pocket volume of both C2A and C2B to understand how all three loops that circumscribe the binding volume respond to the AD3 mutation. We used a convex hull volume as a heuristic to study changes in the volume of the Ca^2+^ binding pocket in the bound and unbound configurations. We defined the volume of the pocket by the convex hull of atoms that coordinated Ca^2+^ using the available structures (C2A:1BYN and C2B:1TJX). The convex hull is defined as the smallest polyhedron that contains each of the coordinating oxygen atoms as vertices (Asp172, Asp178, Asp230, Asp232, and Asp238) in C2A and (Asp303, Asp309, Asp363, Asp365, Asp371) in C2B. From other structures of C2 domains in the presence of Ca^2+^, we expect that the Ca^2+^ -bound volume should be more rigid due to the stabilizing effect of Ca^2+^ [28]. Indeed we observed smaller Ca^2+^ -bound variance from the mean in the binding pocket volume over the course of the simulations, *σ* = 25.39 in C2A^WT^ and *σ* = 9.64 in C2B^WT^ (Table 1., Fig 5) compared to the unbound variance from the mean of *σ* = 66.24 in C2A^WT^ and *σ* = 42.63 in C2B^WT^. In the case of the AD3 mutation, we found that the mean volume and variance of the unbound binding pocket was greater compared to either wild-type volume (Table 1 and Fig 5). Both C2A and C2B increased in volume by approximately 10% (9.9% for C2A and 9.6% for C2B) with the introduction of the AD3 mutation. Interestingly, the volumes of both mutant C2 domains collapsed inward once Ca^2+^ was present and both become exceptionally static throughout the simulation (Table 1 and Fig 5). The volume of the mutant C2A’s Ca^2+^ binding pocket was reduced by 13.8% upon ligand binding; similarly, the mutant C2B’s binding pocket volume was reduced by 13.7% (Table 1).

**Table 1.** Binding pocket volume in reference to Fig 5. Standard Deviation (SD,*σ*) and mean volume are calculated in Ca^2+^-bound and Ca^2+^-unbound C2 domains.

| Ca<sup>2+</sup>-Unbound | Mean vol. (Å<sup>3</sup>) | SD $\sigma$ | Ca<sup>2+</sup>-Bound | Mean vol. (Å<sup>3</sup>) | SD $\sigma$ |
| --- | --- | --- | --- | --- | --- |
| C2A<sup>WT</sup> | 385.67 | 66.24 | C2A<sup>WT</sup> | 134.70 | 25.39 |
| C2A<sup>AD3</sup> | 423.87 | 70.10 | C2A<sup>AD3</sup> | 116.09 | 14.88 |
| C2B<sup>WT</sup> | 193.07 | 42.63 | C2B<sup>WT</sup> | 44.89 | 9.64 |
| C2B<sup>AD3</sup> | 211.70 | 58.00 | C2B<sup>AD3</sup> | 38.73 | 2.41 |

**Figure 5.**
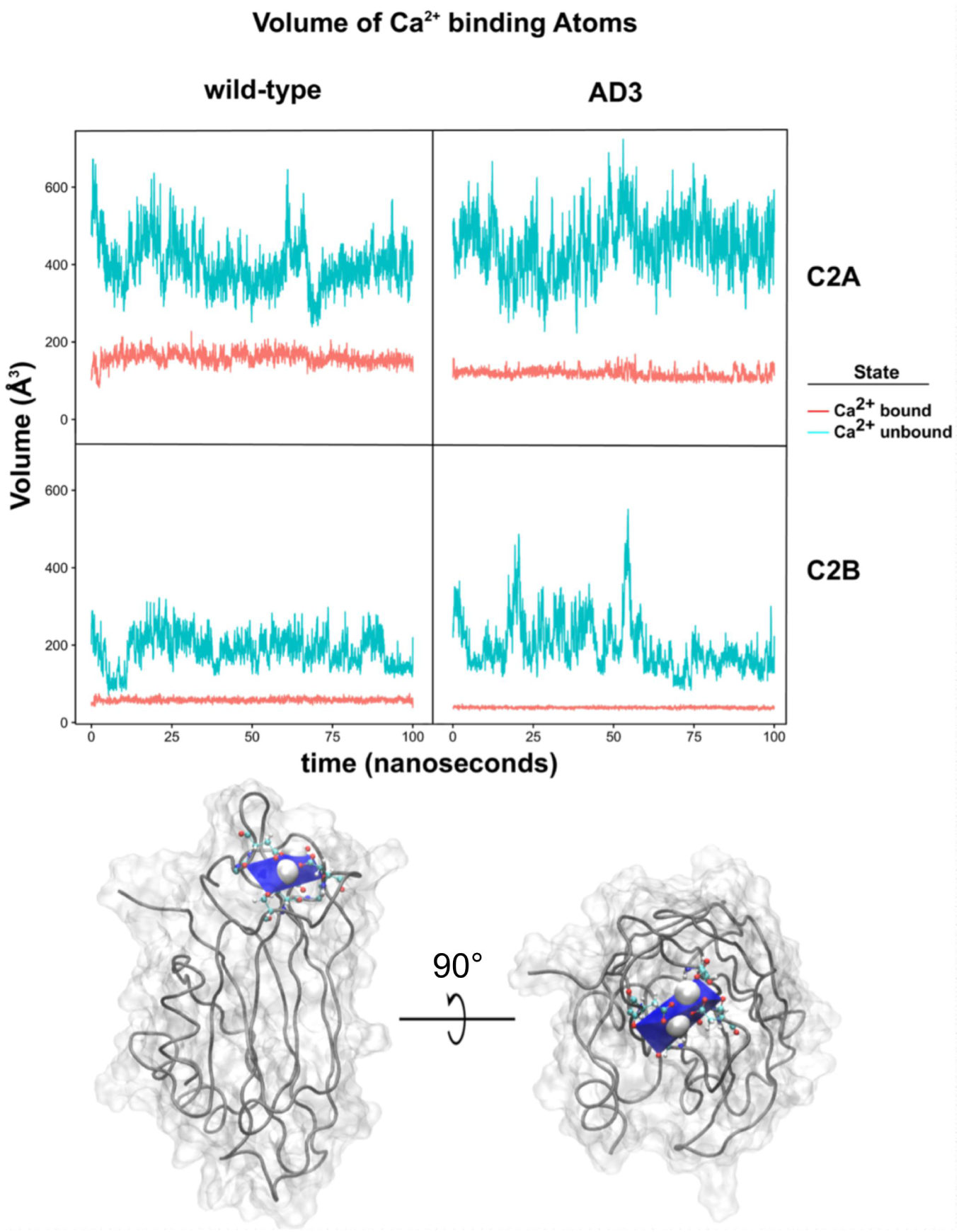
Dynamic volume analysis of binding pocket prescribed by Ca^2+^ coordinating atoms. The volume vs. time of the Ca^2+^ binding pocket is shown in orange; the volume vs time of the unbound volume is shown in cyan. Cartoon representation of the binding volume in blue. Calcium ions are shown as spheres.

### Structural Basis of Dynamic Control

A structural analysis of the interactions involving the AD3 tyrosine H-bond donor in both C2A and C2B shows that a hydrogen bond could link the AD3 residue to the acceptor residue on loop 3. In Syt1 C2A the acceptor would be His-237, and in C2B the acceptor would be Asn-370. Therefore, it is possible that the presence or absence of this H-bond could be the determinant for the activity of loop 3 in the C2 domain. If this is the case, we predict that without the H-bond there should be increased mobility in loop 3 in both C2 domains. Indeed, we observe greater mobility in loop 3 in C2A (Fig 4A). However, unexpectedly, loop 3 of C2B is relatively static throughout the trajectory, while loop 1, on the opposite side of the domain, is more mobile (Fig 4B). These results suggest that the motions of the loops are coupled to the body of the C2 domain.

The variance of the distance between residues measured in Fig 4 is a measure of their coupling. Histogram plots can be used to monitor the dynamic distance between the acceptor residue on loop 3 relative to the absolute position of the AD3 donor residue in wild-type and mutant C2A and C2B (Fig 6 and Table 3). Due to the differences in acceptor amino acids on loop 3 in C2A and C2B, we measured the distances using C*α* atoms instead of the actual H-bonding atoms as these are common distances that can be compared between the C2A and C2B domains. In C2A^WT^ (Fig 6 and Table 3 C2A), there is a prominent unimodal distribution that is the result of the H-bond between His-237 and Tyr-180, which restrains the loop position (Fig 6 and Table 3 C2A, blue histogram). Upon the introduction of the Y180N AD3 mutation, the distance between the donor residue C*α* and the mutant acceptor C*α* becomes shorter as loop 3 collapses away from the Ca^2+^ binding pocket and toward the body of the domain (Fig 6 and Table 3 C2A, orange histogram). The broader distribution of the mutant C2A donor-acceptor distances is primarily due to excessive opening of the Ca^2+^ binding pocket. The bulky tyrosine residue at the AD3 position in many C2 domains seems to block over-opening of loop 3. The lack of the wild-type tyrosine residue in the AD3 mutant domains allows loop 3 almost unrestricted motion. In wild-type C2B, there is also a single distribution of distances between donor and acceptor (Asn370 and Tyr311) suggesting that a similar mechanism operates within C2B. Comparing the variance in backbone distances between C2A and C2B should indicate whether the uncoupling effect of AD3 is more severe in C2A or C2B. The differences between the variances of C2A^WT^ and C2A^AD3^ is (ΔSD) is 0.17; the analogous comparison in C2B is 0.28 (Fig 6 and Table 3). The difference in variance is larger by a factor of 1.6 for C2B than C2A suggesting that a larger ensemble of C2B loop 3 structures is possible relative to C2A. Further, the coupling of loop 3 motions is more pronounced in C2B than in C2A. C2A seems to be regulated by a relatively simple mechanism involving the presence or absence of the H-bond between His-237 and Tyr-180. C2B appears to be more complicated.

**Table 3.**
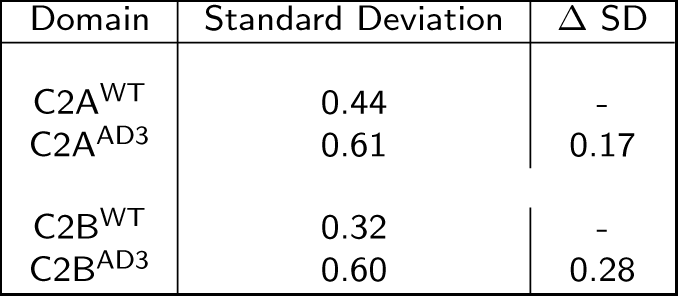
Width of the distance distribution between C*α*-180 and C*α*-237 in C2A, and C*α*-311 and C*α*-370 in C2B. The AD3 locus is C*α*-237 in C2A and C*α*-370 in C2B.

**Figure 6.**
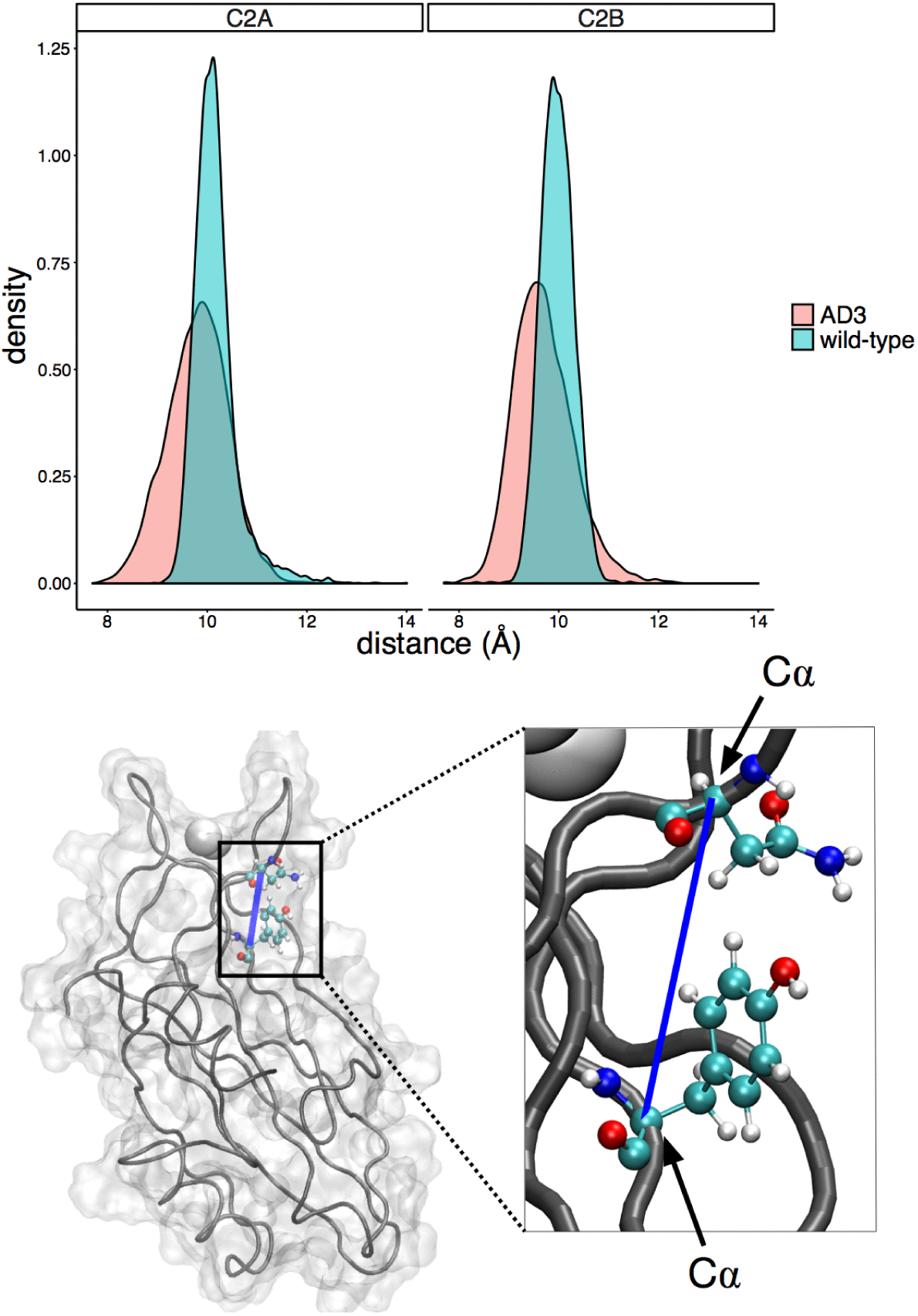
Histogram of donor:acceptor C*α* distances in C2A and C2B in wild-type and the AD3 mutant. The bottom structure is a Connelly surface representation illustrating the distance measured between C*α* atoms of Asn-370 and Tyr-311 in Syt1 C2B. A zoom out of the actual distance is shown as a ball-and-stick representation to the right of the Connelly surface. The distance between C*α* atoms is highlighted as a blue line

To investigate the mechanism for control of loop dynamics in C2B, we used correlation network analysis to examine the differences in intra-*β*-sheet connectivity between the two *β*-sheets of the C2B domain. The correlated motion network of C2B^WT^ reveals that loop 1’s motion is correlated with the motion of C*α* atoms in main body of the C2B domain, but not loop 3 (Fig 7A). In C2B^AD3^, however, the correlated motions of loops 1 and loop 3 fully decouple from the C2B domain (Fig 7).

**Figure 7.**
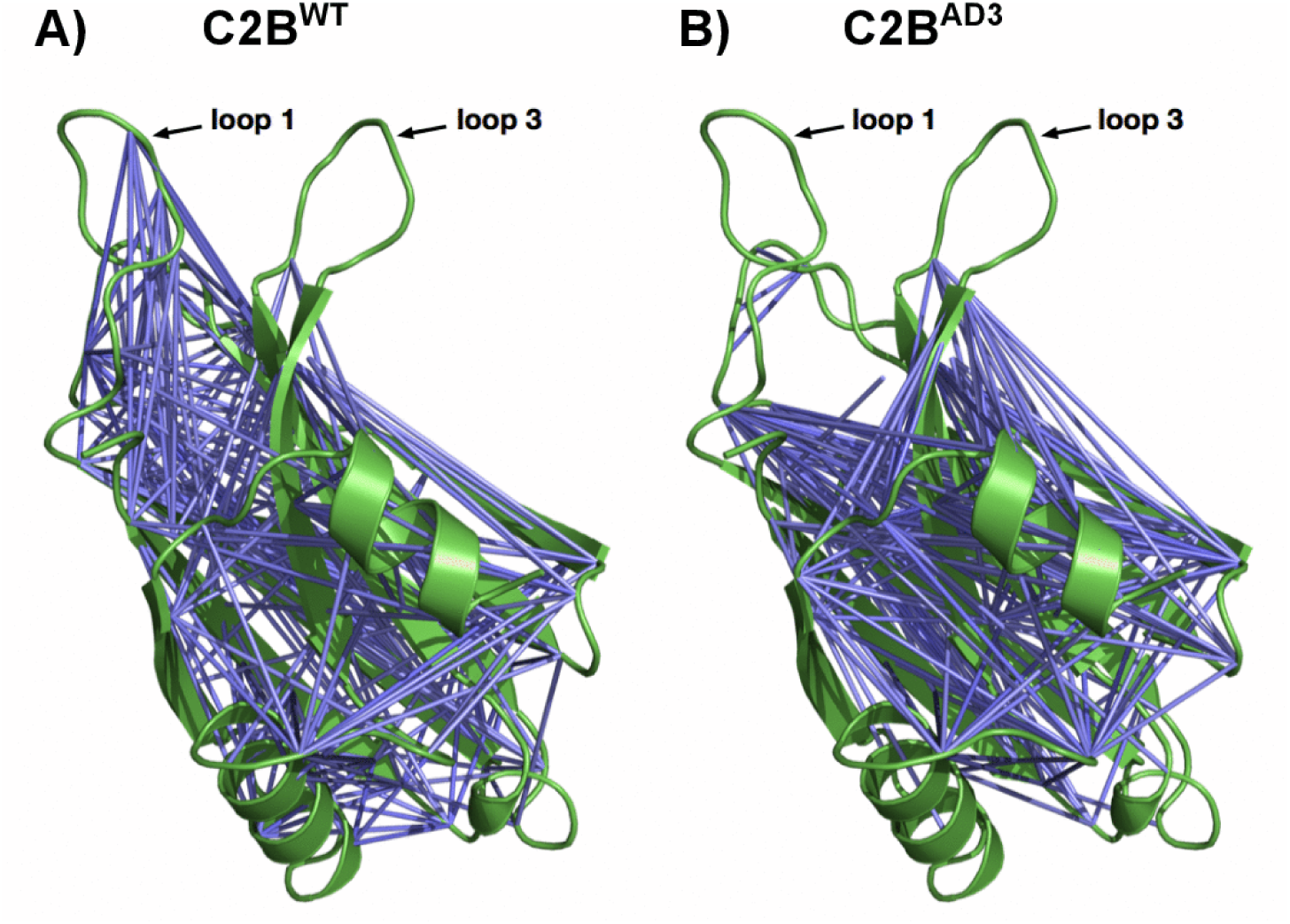
Dynamic network analysis of C2B. Blue bars indicated that C*α* motions are correlated above the threshold of 0.4. A) Correlated Network Analysis for Syt1 C2AWT. B) Correlated Network Analysis for Syt1 C2B^AD3^.

## Discussion

Once an action potential invades a neuron, Ca^2+^ enters the cell through voltage-gated Ca^2+^ channels that are located near the active zone [31]. Ca^2+^ -bound Syt1 can then associate with acidic phospholipids and the SNARE complex [32], with the end result of neurotransmitter release to the synaptic cleft. Syt1 is, therefore, at the crux of the exocytosis signaling network, and has been identified as the Ca^2+^ sensor that triggers fast, synchronous neurotransmitter release in neurons [33,34].

The biophysical properties of Syt1 are well-suited for Ca^2+^ sensing in the neuron. Syt1 coordinates calcium ions in a solvent-exposed pocket at the apex of each C2 domain [14,35]. The binding pocket for Ca^2+^ is formed by three loops; however, only loops 1 and loop 3 contribute acidic residues that coordinate calcium ion. This solvent-exposed, jaws-like architecture lends itself to a relatively low Ca^2+^ binding affinity [36], making Syt1 the ideal trigger to respond to rapid changes in the calcium ion concentration at the active zone. Despite the progress made in understanding the biophysics of exocytosis at atomic detail, questions remain concerning how C2 domains function within the limitations of the transient Ca^2+^ gradients present in the neuron. By examining the consequences of mutations at the AD3 locus in the Syt1 C2 domains using molecular dynamics techniques, we can understand how the modulation of loop mobility contributes to the activity of synaptotagmins in cells.

The AD3 locus of synaptotagmin is interesting for at least two reasons. First, the locus is not directly associated with the Ca^2+^ binding site of C2 domains, yet it affects the ability of the domain to respond to Ca^2+^ [23], as electrophysiology experiments using neurons from AD3 flies showed a clear deficit in Ca^2+^ -evoked release [33]. Indeed, we have shown that in the minimal fusion assay, the AD3 mutation in Syt1 reduces the ability of SNARE-laden vesicles to fuse (Fig 2). Second, the AD3 point mutation is located within the highly conserved’ SDP<u>Y</u>VK’ sequence that occurs in many C2 domains (Fig 1), not only synaptotagmins. The question then becomes” How does AD3 alter Syt1 function?”, as there must be a physical link between the AD3 locus and the Ca^2+^ -binding apparatus of the C2 domain.

In the wild-type C2A domain, both loops 1 and 3 exhibit mobile behavior without Ca^2+^ (Fig 3). Indeed, the temperature factors of loop 1 in most C2A X-ray structures tend to be higher than average, indicating disorder of that loop [14]. By plotting the RMSF values of wild-type versus AD3 mutant trajectories (Fig 4A and 4B), we see that there is additional mobility contributed by the AD3 point mutation on loops 1 and 3; loop 3 is particularly interesting. In the wild-type C2A domain, the conserved Tyr residue likely forms an H-bond to a residue on loop 3 of the Ca^2+^ binding loop, thus restricting the ensemble of possible structures for loop 3. Our simulations confirm that when the AD3 locus is mutated to Asn, loop 3 decouples and becomes more flexible relative to the wild-type C2A domain (Fig 4A). On average the increase in loop flexibility from the AD3 mutation in C2A adds an additional 16 Å^3^ of volume to the binding pocket of C2A. The additional volume in C2A could potentially accommodate at least two additional calcium ions. The mean volume of C2B is 9.2% larger with the AD3 mutation; this additional volume in C2B could also hold two more calcium ions.

C2A and C2B of synaptotagmin are clearly homologous domains, so it is reasonable to expect two similar domains to function similarly with respect to Ca^2+^. Therefore, we would guess that C2B would also behave like C2A when the conserved tyrosine residue is mutated to an asparagine. However, this does not appear to be the case. Loop 3 in C2B becomes relatively more rigid and loop 1 becomes more flexible. The net result is still an overall increase in the net volume of the Ca^2+^ binding pocket, but by different mechanisms. In Fig 3, each of the three of the Ca^2+^ -binding loops in C2B has different flexibility throughout the simulations relative to C2A. Further, the iso-dynamic comparison of wild-type versus AD3 C2B shows that loop 3 is essentially rigid in C2B; it is loop 1 that is mobile in this case. That is unusual because there is no simple physical connection between the AD3 locus and loop 1 in C2B. The volume analysis is more telling. The wild-type volume of the Ca^2+^ binding pocket of C2B is smaller than C2A. Once Ca^2+^ binds to C2B, the inward loop collapse is more pronounced. In the context of the AD3 mutation, the average volume of C2B is larger than that of the wild-type protein; however, when Ca^2+^ binds, loops 1 and 3 collapse around the cations with an exceptionally small volume and a very small variance. There is a clear impact in the interaction of the AD3 locus, the dynamics of loop3 and the shape of the Ca^2+^ binding pocket of C2A, but the question remains as to where is the linkage between Tyr-311 and loop 1 in C2B.

Another computational method to assess the AD3 phenotype is through correlated-motion analysis. Correlated motion is a property of *β*-sheet proteins where the dynamics of the *β*-strands are coupled by the hydrogen-bonding network that stabilizes the sheet [37]. The linkage includes not only backbone H-bonds but side chains that contribute H-bonds. An analysis of the H-bonds formed throughout the time course of the trajectories in C2A and C2B, whether wild-type or mutant, bound or unbound, shows that C2B possesses, on average, 25 more H-bonds than C2A (Table 2). The H-bond analysis includes all secondary structure in C2A and C2B. Due to the clamshell-like *β*-sandwich construction of C2 domains, there is no continuous H-bond path that connects the two *β*-sheets. The most apparent path of H-bond connectivity exists at the loops between *β*-sheets with loop 2 being the most prominent intermediary in C2B. As C2B has inherently more H-bonds, the H-bonds could be utilized to propagate information across C2B. Therefore, the interaction between Tyr-311 and loop 1 could be mediated through inter-sheet H-bonding that communicates using backbone interactions influenced by Tyr-311 through loop 2 and finally up to loop 1, as there is additional flexibility observed in loop 2 that is not reflected in the trajectories of C2A (Fig4 C2A and Fig 4 C2B).

**Table 2.** Number of H-bonds in C2 domains throughout the course of the simulations. The value in parenthesis refers to the number of H-bonds formed while the domain was occupied with two calcium ions.

| Domain | Mean number of H-bonds |
| --- | --- |
| C2A<sup>WT</sup> (Ca<sup>2+</sup>) | 61.50 (62.07) |
| C2A<sup>AD3</sup> (Ca<sup>2+</sup>) | 60.72 (64.35) |
| C2B<sup>WT</sup> (Ca<sup>2+</sup>) | 86.25 (86.88) |
| C2B<sup>AD3</sup> (Ca<sup>2+</sup>) | 85.85 (86.94) |

Calcium ions are known to stabilize the loops in C2 domains [28,27]. In our simulations of unbound C2A, as shown in Table 1, the accumulated trajectories of C2A have a standard deviation of *σ* = 66.24 over the 100 ns time course. When Ca^2+^ is included in the simulation, the standard deviation of the volume of the Ca^2+^ binding pocket shrinks considerably for both domains (Table 1). Interestingly, C2A with the AD3 mutation and bound Ca^2+^ has a mean volume of 116 Å^3^, which is less than wild-type. The collapse of this binding pocket is likely the result of the volume being dominated solely by the bridging of loops 1 and 3 by Ca^2+^. Therefore, in addition to preventing over-opening of the Ca^2+^ binding pocket, the same mechanism may prevent over-collapse. The importance of the AD3 regulatory mechanism is evident in the C2A-C2B tandem structure. In this crystal structure, the proximity of the C2B domain formed an H-bond between His-237 in C2A and Thr-406 in C2B [10]. The decoupling of His-237 from Tyr-180, in the presence of C2B, resulted in the collapse of loop 3 and concomitant distortion of the Ca^2+^ binding pocket of C2A in that particular domain [10]. The AD3 mutation in the isolated C2A domain is mimicked in the C2A domain of the C2A-C2B X-ray crystal structure [10].

The two tandem C2 domains of synaptotagmin are not simply two duplicated domains with redundant activities. Recent data indicate that the two domains co-operate, each with unique contributions, to accomplish the task of membrane fusion [38, 39, 40, 41, 10]. As complementary protein domains, both C2A and C2B have unique biophysical properties with respect to Ca^2+^ binding and phospholipid bi-layer modification properties [42,43]; further, both domains respond differently to the AD3 point mutation. In the unbound state of C2A, the hydrogen bond between Tyr-180 and His-234 that usually maintains an open and relatively fixed volume is lost in C2A^AD3^ (Fig 6 and Table 3), and the flexibility of loop 3 increased accordingly (Fig 4 C2A). The increased flexibility of loop 3 results in an increase in the volume of the Ca^2+^ -binding pocket relative to C2A^WT^. Since the Ca^2+^ -binding loops and lipid binding activity are linked in C2 domains, the abilities of Syt1 C2AB to bind Ca^2+^, to engage SNAREs and to fuse vesicles and are also affected (Fig 2A, 2B, and 2C). In the C2B domain, where the AD3 mutation was originally identified, loop 3 is relatively static compared to C2A; however, the dynamic motion of loop 1 is decoupled from the rest of the domain in the mutant domain. The decoupled motion in loop 1 could be associated with cooperative inter-strand dynamics that link from the site of the mutation to loop 1 (Fig 7). C2A^WT^ is therefore regulated primarily by a doorstop-like mechanism, where loop 3 is blocked from either over-opening or over-collapsing by the interaction between His-237 and Tyr-180. C2B appears to be regulated not only by the presence of the Tyr-311 residue acting as a steric block to help shape the Ca^2+^ binding pocket, but also the H-bond network throughout the C2B domain facilitates communication between Ca^2+^ -binding loops and the body of the C2 domain. Hence, the original AD3 mutation, C2B^AD3^, first described by Diantonio *et al.* [20] is likely the result of uncoupling an innate regulatory mechanism present within C2 domains that regulates the volume and shape, and therefore the Ca^2+^ -binding affinity, of the Ca^2+^ -binding pocket.

## Materials and Methods

### Recombinant Proteins and Protein Purification

cDNA encoding the cytoplasmic domain of Syt1 (denoted C2AB, amino acids 96-421) was provided by T.C. Südhof (Stanford, CA); a glycine residue was substituted to correct the D374 mutation [44]. cDNA encoding synaptobrevin-2 (syb) and syntaxin-1A (syx) were provided by J.E. Rothman (New Haven, CT). cDNA encoding SNAP-25B was provided by M.C. Wilson (Albuquerque, NM). The AD3 mutant form of rat Syt1 C2B (Y311N) was generated using QuikChange Site-Directed Mutagenesis Kit (Stratagene). Syt1 C2AB and the AD3 mutant were subcloned into pGEX-4T vectors and expressed as GST-tagged fusion proteins. Proteins were purified using glutathione–Sepharose beads (GE Healthcare) and cleaved with thrombin to remove the GST-tag, as described [45]. All three full-length SNARE proteins were individually subcloned into a pTrcHis vector and expressed as His6-tagged fusion proteins; SNAP-25B and syntaxin 1A were also subcloned into a pRSF-Duet vector. Proteins were purified using Ni-Sepharose beads (GE Healthcare), as previously described [45].

### Phospholipids

Synthetic lipids were purchased from Avanti Polar Lipids: palmitoyl-2-oleoyl-sn-glycero-3-phosphocholine (phosphatidylcholine, PC), 1,2-dioleoyl-sn-glycero-3-phospho-l-serine (phosphatidylserine, PS), 1-palmitoyl-2-oleoyl-sn-glycero-3-phosphoethanolamine (phosphatidylethanolamine, PE),1,2-dipalmitoyl-sn-glycero-3-phospho-ethanolamine-N-(7-nitro-2-1,3-benzoxadiazol-4-yl) (NBD-PE), N-(lissamine rhodamine B sulfonyl)-1,2-dipalmitoyl-sn-glycero-3-phosphoethanolamine (Rho-PE).

### *In vitro* Fusion Assay

For *in vitro* fusion assays, SNARE-bearing vesicles were prepared as previously described [6]. The lipid compositions for vesicles were as follows: 15% PS, 27% PE, 55% PC, 1.5% NBD-PE, and 1.5% Rho-PE for v-SNARE vesicles, 15% PS, 30% PE, and 55% PC for t-SNARE heterodimer and 25% PS, 30% PE, and 55% PC for syntaxin-only vesicles. Fusion between v-SNARE vesicles and t-SNARE heterodimer, or syntaxin-only, vesicles was monitored using a Synergy HT multi-detection microplate reader (Bio-Tek). For assays, the cytoplasmic domain (C2AB) of Syt1 or the AD3 mutant (1 *µ*M) was mixed with 0.2 mM EGTA, 0.5 *µ*l v-SNARE vesicles, and 4.5 *µ*l t-SNARE vesicles for standard assays or 4.5 *µ*l syntaxin-only vesicles plus 3 *µ*M soluble SNAP-25 for split t-SNARE assays. During the experimental run, we added Ca^2+^ (to a 1 mM final concentration), and the reaction was monitored for an additional 60 min. To give the maximum fluorescence signal, detergent was added after each experiment; this value was used for normalization. In addition to representative traces, the extent of fusion (t = 80 min) was plotted for each protein.

## Molecular Dynamics

We obtained the crystal structures of *Rattus norvegicus* Syt1 C2A (4WEE) and Syt1 C2B (1TJX) [46] from the Protein Data Bank. We then generated models of the representative mutations in each C2 domain using the mutagenesis wizard in Pymol [29]. Autopsf, a TCL utility, was used to generate the PSF file from the oriented PDB file. The structures were solvated using the solvate TCL utility in a cube of approximately 7,000 water molecules (TIP3) with a positive and negative padding of 10 Åon each axis. Residue 226 in C2A and residue 359 in C2B were restrained with harmonic restraints to dampen coordinate drift within the simulation box. Each structure was simulated using NAMD [47] with different random seeds for 50,000,000 frames with a time step of two femtoseconds and a write frequency of 25,000. Periodic boundary conditions were used with PME and a grid spacing of 1.0. We used rigid bonds and constant group pressure control. The pressure was controlled with a Langevin piston set at 1.01325 bar (atmospheric pressure). The temperature was controlled at 310 K with Langevin dynamics, and a dampening coefficient of 5 *ps*^-1^ was applied to each trajectory. We validated that the trajectories reached equilibrium by calculating the root mean squared displacements of all atoms in the protein after alignment over the course of the run. Each unique simulation setup was run three times for a total of 300 ns of simulation time. The simulation time we used is consistent with other recent MD studies of synaptotagmin [48,49].

To study the differences between the C2 domains of wild-type the AD3 mutants in both the Ca^2+^ -bound and unbound states, we ran several simulations of Ca^2+^ - bound *Rattus norvegicus* Syt1 1BYN (C2A) [50] and 1TJX (C2B) [46]. In the cases where Ca^2+^ -bound domains were required for analysis, the cations were coincident with known positions within the domains and coordination was consistent with known X-ray [46] and NMR structures [50]. Molecular dynamics simulations were run as above while holding the Ca^2+^ at its observed location within the binding pocket. We also restrained a single C*α* at the center of the protein to dampen coordinate drift. To reach relative equilibrium, we selected a Ca^2+^ -bound structure for simulation restart after the initial restrained run. We also used a single Ca^2+^ - bound frame, excluding the calcium ion, as a starting position for several runs both with and without exogenous Ca^2+^ in 150 mM K^+^ and Cl^-^. These 100 ns simulations were then used for structural analysis including the RMSF, volume, and distance analyses.

## Iso-dynamic Plot Analysis

To compare the relative mobility of the AD3 mutant and wild type, we developed a method to compare and quantify differences in RMSF between groups of residues (Fig 4A and 4B). Each point in the plane represents an amino acid with its *x* component corresponding to the wild type RMSF and the *y* corresponding to AD3 RMSF. A point in the plane (*a, b*) with *a* = *b* has equal RMSF in wild type and AD3, and such points will lie on the iso-dynamic line *y* = *x*. Residues which are affected by the mutation will have a larger residual from the line. Further, a point (*a, b*) with *b > a* that lies above the iso-dynamic line will have a greater mobility in the AD3 mutant, and the reverse holds for points below the line. We can quantify the effect of the AD3 mutation on a residue’s mobility by measuring its distance from the *y* = *x* iso-dynamic line.

## Molecular Dynamics Analysis

Molecular dynamics trajectories were analyzed using MDAnalysis [51], SciPy [52], Bio3d [53], and VMD [54]. The hull calculations were performed using the scipy.spatial.ConvexHull [52] implementation of the Quickhull algorithm [55].

## Correlation Network Analysis

Network analysis of correlated motions was used to identify aspects of the C2B domain with linked motion [56]. A weighted network graph was computed where each node represents a residue and weights of the connections between nodes represents the respective cross-correlation of their displacements [57]. The network edges that we used were constructed based on the C*α*-C*α* cross-correlation value between all residues across all replicate simulations in the respective domains. Network edges were added for residue pairs with correlations ≥ 0.4 in all simulations and with a C*α*-C*α* distance ≤ 10 Å. The Bio3d package was used to compute all correlation network analyses [53].

## Declarations

### Acknowledgements

The authors acknowledge the Texas Advanced Computing Center (TACC, URL: http://www.tacc.utexas.edu) at The University of Texas at Austin and the High-Performance Computing Center (HPCC, http://cmsdev.ttu.edu/hpcc) at Texas Tech University at Lubbock for providing high performance computing resources that have contributed to the research results reported in this paper. NAMD was developed by the Theoretical and Computational Biophysics Group in the Beckman Institute for Advanced Science and Technology at the University of Illinois at Urbana-Champaign.

### Funding

Research reported in this publication was supported by National Institute of Arthritis and Musculoskeletal and Skin Diseases of the National Institutes of Health under award number AR063634 to RBS and MH061876 to ERC. The funders had no role in study design, data collection and analysis, decision to publish, or preparation of the manuscript.

### Availability of data and materials

The data and scripts supporting the claims in this paper will be available on the following link: https://goo.gl/Gkb6ao

### Authors’ contributions

RBS, AGM, and PJR conceived the study and designed the experiments. PJR performed all computational experiments and data analysis. RBS secured funding for the work. RBS wrote the manuscript. RBS and PJR generated the figures. All authors have read and approved the manuscript.

### Ethics approval and consent to participate

Not applicable

### Consent for publication

Not Applicable.

### Competing interests

The authors declare that they have no competing interests.

